# EZH2 represses mesenchymal genes and upholds the epithelial state of breast carcinoma cells

**DOI:** 10.1101/2023.03.13.532335

**Authors:** Amador Gallardo, Lourdes López-Onieva, Efres Belmonte-Reche, Iván Fernández-Rengel, Andrea Serrano-Prados, Aldara Molina, Antonio Sánchez-Pozo, David Landeira

**Author notes:** **Correspondence** to David Landeira.

## Abstract

Emerging studies support that the Polycomb Repressive Complex 2 (PRC2) regulates phenotypic changes of carcinoma cells by modulating their shifts among metastable states within the epithelial and mesenchymal spectrum. This new role of PRC2 in cancer has been recently proposed to stem from the ability of its catalytic subunit EZH2 to bind and modulate the transcription of mesenchymal genes during epithelial-mesenchymal transition (EMT) in lung cancer cells. Here, we asked whether this mechanism is conserved across different types of carcinomas. By combining TGF-β-mediate reversible induction of epithelial to mesenchymal transition and pharmacological inhibition of EZH2 activity we demonstrate that EZH2 represses a large set of mesenchymal genes and favours the residence of breast cancer cells towards the more epithelial spectrum during EMT. In agreement, analysis of human patient samples support that EZH2 is required to efficiently repress mesenchymal genes in breast cancer tumours. Our results indicate that PRC2 operates through similar mechanisms in breast and lung cancer cells. We propose that PRC2-mediated direct transcriptional modulation of the mesenchymal gene expression program is a conserved molecular mechanism underlying cell dissemination across human carcinomas.

## Introduction

Polycomb group (PcG) proteins are hallmark epigenetic regulators of embryo development, stem cell differentiation and cancer (Blackledge & Klose, 2021; Piunti & Shilatifard, 2021; Schuettengruber *et al*, 2017). PcG proteins associate to form multimeric complexes termed Polycomb Repressive Complexes 1 and 2 (PRC1 and PRC2) that can post-translationally modify histone tails and repress gene transcription. RING1A/B is the catalytic subunit of PRC1 and monoubiquitinates lysine 119 on histone H2A (H2AK119ub). Likewise, EZH1/2 harbours the catalytic activity of PRC2 and trimethylates lysine 27 on histone H3 (H3K27me3). The coordinated activity of PRCs leads to the formation of chromatin domains enriched in H2AK119ub and H3K27me3 that facilitate transcriptional repression of hundreds of genes across the genome in a cell-type-specific manner. In stem cells, the activity of PRCs lead to the transcriptional repression of hundreds of lineage-inappropriate genes, contributing to maintain a specific gene expression program and define stem cell identity (Blackledge & Klose, 2021; Piunti & Shilatifard, 2021; Schuettengruber *et al*., 2017).

In the context of cancer, initial studies revealed that PcG proteins act as oncogenes through the transcriptional repression of the *INK4A/ARF* (*CDKN2A*) tumour suppressor locus (Bracken *et al*, 2007; Jacobs *et al*, 1999). However, subsequent studies confirmed that this was just one aspect of the role of PRC in cancer, because different PRC subunits can have senescence-independent prooncogenic activity or tumour suppressor function (Piunti & Shilatifard, 2021; Serresi *et al*, 2016). In fitting with its role as a regulator of cell identity in stem cell biology, recent studies support that PRC2 can regulate dynamic phenotypic changes of cancer cells by modulating their transition between metastable states within the epithelial and mesenchymal spectrum through poorly understood mechanisms (Gallardo *et al*, 2022; Serresi *et al*., 2016; Serresi *et al*, 2018; Zhang *et al*, 2022b). Reversible transition between the epithelial and mesenchymal states vertebrate carcinoma cell dissemination (Lambert *et al*, 2017), and inhibitors of the PRC2 catalytic subunits EZH2 are currently being developed to treat several types of carcinomas as main or adjuvant therapy (Huang *et al*, 2022; Kim & Roberts, 2016). Therefore, understanding the molecular basis of the function of PRC2 during EMT is crucial for a precise understanding of the molecular basis of cancer dissemination, and for successful application of PRC inhibitors in the clinics.

In breast cancer, comprehensive evidence supports that EZH2 facilitates metastasis (Hirukawa *et al*, 2018; Moore *et al*, 2013; Yomtoubian *et al*, 2020; Zhang *et al*, 2022a). However, the underlying molecular mechanism is unclear. Initial reports suggested that EZH2 might impact metastasis progression through non-canonical pathways independent of H3K27 methyltransferase activity (ie. p38 signalling and integrin B1-FAK) (Moore *et al*., 2013; Zhang *et al*., 2022a). Other studies highlight that EZH2 regulates repression of specific genes through H3K27me3 (ie. FOXC1 or GATA3) (Hirukawa *et al*., 2018; Yomtoubian *et al*., 2020). Interestingly, a recent report supports that the regulation of metastasis by PRC2 might be linked to the modulation of EMT (Zhang *et al*., 2022b). This is in consonance with recent findings in lung cancer where it has been shown that loss of function of EZH2 leads to acquisition of mesenchymal features and changes in tumour colonization capacity (Gallardo *et al*., 2022; Serresi *et al*., 2016; Serresi *et al*, 2021; Serresi *et al*., 2018; Zhang *et al*., 2022b). This effect has been proposed to stem from the ability of EZH2 to directly bind the gene promoter regions and co-ordinately modulate the transcription of the mesenchymal gene expression program through H3K27me3 during EMT (Gallardo *et al*., 2022). Here, we asked whether transcriptional regulation of mesenchymal genes by EZH2 also occurs in breast carcinoma cells. We found that EZH2 represses a large set of mesenchymal genes and promotes the residence of breast cancer cells in a more epithelial state. We propose that direct transcriptional modulation of the mesenchymal gene expression program by PRC2 is a conserved molecular mechanism across different types of carcinomas that contributes to endorse cells with the plasticity required for efficient cell dissemination.

## Results

### EZH2 represses mesenchymal genes in breast carcinoma cells

To analyse the function of EZH2 in breast cancer we focused on the human MCF-7 cell line because it is a well-stablished system to study the molecular basis of metastasis in breast adenocarcinomas. These cells are homozygous null mutant for the *CDKN2A* locus, which makes them a good model to study the *CDKN2A*-independent function of EZH2 in cancer. We first set to identify what genes are directly regulated by EZH2 in MCF-7 cells. EZH2 catalyses H3K27me3 at the promoter regions of target genes, and thus, the genome-wide distribution of H3K27me3 is a surrogate measure of EZH2 binding (Blackledge & Klose, 2021; Comet *et al*, 2016). Analysis of H3K27me3 enrichment maps revealed that EZH2 target the promoter region of 1954 genes in MCF-7 cells (Figure 1A, Table S1). In pluripotent cells, many H3K27me3-repressed target genes display chromatin features of active transcription such as trimethylation of H3K4 (H3K4me3) and they are usually referred to as bivalent genes (Macrae *et al*, 2022). Bivalent chromatin seems to facilitate their transcriptional activation during lineage transition (Macrae *et al*., 2022). Interestingly, comparison of H3K27me3 and H3K4me3 genome-wide binding maps in MCF-7 cells revealed that 1141 genes (58.4%) out of the 1954 genes marked by H3K27me3 displayed a bivalent state, and hence they accumulated both modifications at their promoter region (Figure 1A,B, Table S1). Importantly, bivalent genes were enriched for genes involved in the regulation of mesenchymal features, EMT and cell migration, and included genes that are widely used as mesenchymal markers such as *N-CADHERIN* and *SNAI2* (Figure 1C, 1D, S1A). Thus, we concluded that in MCF-7 cells, EZH2 binds a large set of mesenchymal genes that display features of bivalent chromatin. We hypothesized that this bivalent state might facilitate the coordinated transcriptional activation of mesenchymal genes in response to signalling molecules during EMT.

**Figure 1.**
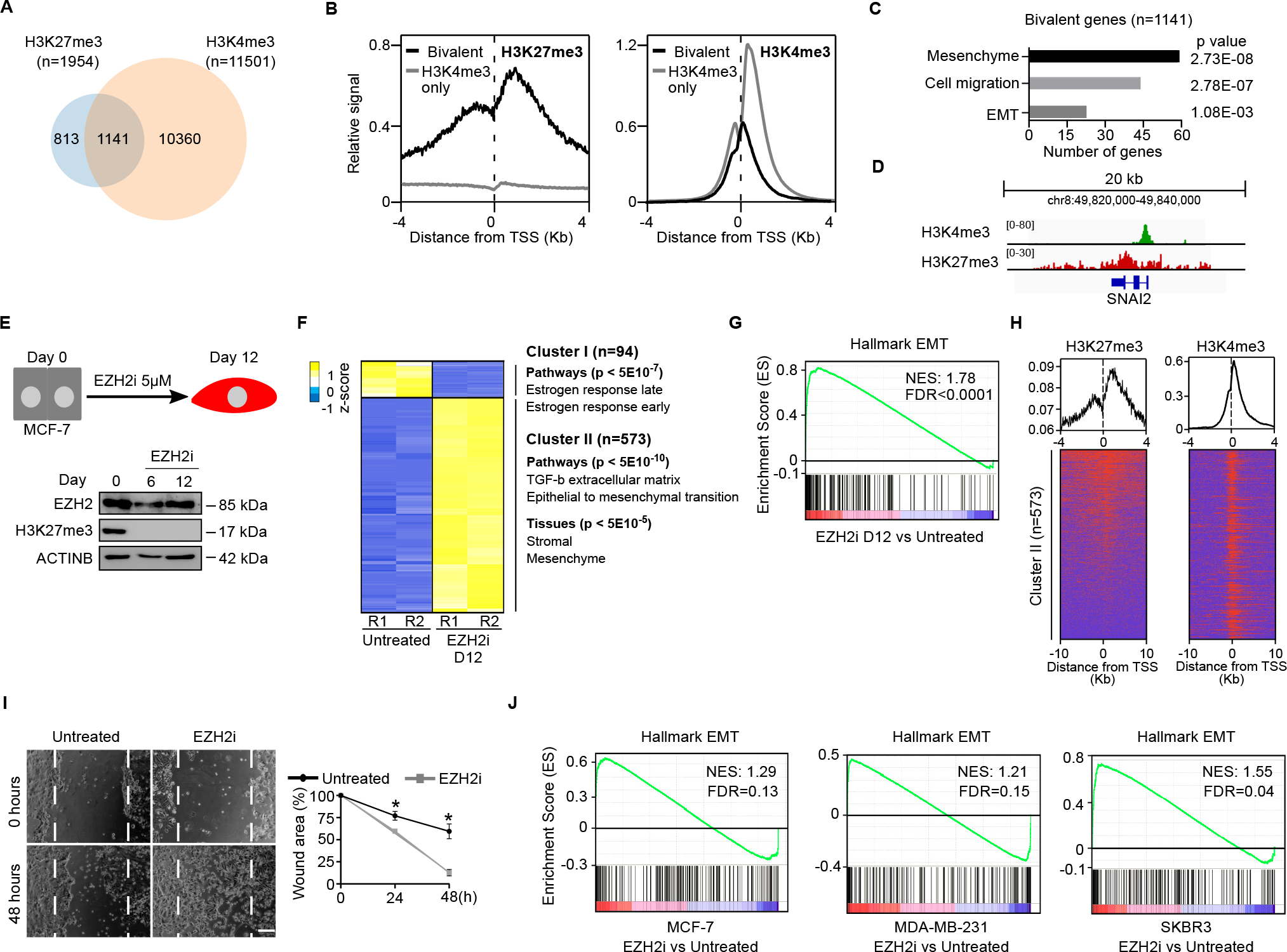
EZH2 represses mesenchymal genes in breast carcinoma cells. **(A)** Venn diagram comparing sets of gene promoters displaying enrichment of H3K27me3 or H3K4me3 at their promoter regions by ChIP-seq in MCF-7 cells. **(B)** Plots showing the ChIP-seq average binding profile of H3K27me3 and H3K4me3 around the TSS of gene promoters identified in (A) as bivalent (H3K27me3+H3K4me3) or H3K4m3-only. **(C)** Gene Ontology analysis of bivalent genes identified in (A). **(D)** Genome browser view of H3K4me3 and H3K27me3 binding profiles at the *SNAI2* locus in MCF-7 cells. **(E)** Schematic diagram of the treatment of MCF-7 cells with EZH2i (upper panel). Lower panel shows western blot analysis of whole-cell extracts comparing the levels of EZH2 and H3K27me3 during the experiment. ACTIN B provides a loading control. **(F)** Heatmap analysis of mRNA expression of 667 differentially expressed genes upon 12 days of EZH2i treatment (FC > 4, p < 0.05) by RNA-seq of two independent replicates (R1 and R2) in MCF-7 cells. **(G)** GSEA of EMT associated genes in MCF-7 cells treated compared to untreated with EZH2i for 12 days. Normalized Enrichment Score (NES) and false discovery rate (FDR) are indicated. **(H)** Plots showing H3K27me3 and H3K4me3 enrichment measured by ChIP-seq around the TSS of the promoters of 573 genes identified in cluster II in figure 1F. **(I)** Brightfield images and quantification of cultured wound healing assays using MCF-7 cells that have been treated or not with EZH2i for four days. Mean and SEM of 3 experiments are shown. Asterisks indicate statistical significance using a Mann-Whitney test (* p<0.05). **(J)** GSEA of EMT associated genes in MCF-7, MDA-MB-231 and SKBR3 cells plated at low density and treated with EZH2i for 14 days, compared to untreated controls. Normalized Enrichment Score (NES) and statistically significant false discovery rate (FDR<0.25) are indicated.

To address whether EZH2 represses the mesenchymal gene expression program through H3K27me3 in breast cancer cells we treated MCF-7 cells with a highly specific small molecule that inhibits EZH2 methyltransferase activity (GSK126, thereafter referred to as EZH2i) (McCabe *et al*, 2012) and analyse whether the loss of H3K27me3 induced the activation of the mesenchymal gene expression program. Inhibition of EZH2 led to a drastic reduction of global levels of H3K27me3 without sensibly affecting EZH2 protein stability (Figure 1E) and produced only mild inhibition of cell proliferation that did not impair long-term culture of MCF-7 cells (Figure S1B). Importantly, reduced levels of H3K27me3 during twelve days of culture led to a robust transcriptional activation of the mesenchymal marker *N-CADHERIN* (Figure S1C), which is a H3K27me3-positive direct target of EZH2 (Figure S1A). Likewise, analysis of the expression of master EMT transcription factors (EMT-TFs) revealed an evident specific activation of the *SNAI2* gene (Figure S1D and S1E), which is also enriched for H3K27me3 and H3K4me3 at its promoter region (Figure 1D). As expected, activation of mesenchymal genes upon EZH2 inhibition led to downregulation of epithelial marker *E-CADHERIN* (Figure S1F and S1G). Importantly, transcriptome profiling analysis by mRNA sequencing (mRNA-seq) demonstrated that treatment of MCF-7 cells with EZH2i induced a global reorganization of the transcriptional program that involved the upregulation of 573 genes enriched in EMT pathways (cluster II in Figure 1F) and mesenchymal markers (Figure 1G). As expected, many of these genes displayed H3K27me3 and H3K4me3 at their promoter regions (Figure 1H). In addition, we noticed that genes involved in estrogen response were downregulated upon EZH2i-treatment (Figure 1F, cluster I), suggesting that activation of EMT induces the inactivation of the estrogen pathway signalling in MCF-7 cells. Importantly, activation of mesenchymal genes in MCF-7 cells upon depletion of H3K27me3 was functionally relevant because cells treated with EZH2i displayed obvious increased mobility compared to untreated cells in cultured wound healing assay (Figure 1I). Thus, we concluded that EZH2 targets and represses the transcription of a large set of bivalent mesenchymal genes in the breast cancer cell line MCF-7.

To confirm that EZH2 functions as a gatekeeper of epithelial identity in breast carcinoma cells we analysed whether inhibition of EZH2 led to the transcriptional activation of mesenchymal genes in cell lines derived from different types of metastatic breast adenocarcinomas: MCF-7 cells (ER+/PR+-/Her2-), SKBR3 cells (ER-, PR-, Her2+) and MDA-MB-231 cells (ER-/PR-/Her2-). Cells were treated with EZH2i for four days and then plated at low density to form colonies during fourteen extra days in the presence of EZH2i. Inhibition of EZH2 induced minor effects on cell growth (Figure S1H), confirming the suitability of these cell lines to study the proliferation-independent role of EZH2. EZH2i-treated cells displayed unpacked colonies composed by cells with more elongated and fusiform morphology typically associated with the mesenchymal state, as compared to non-treated control cultures composed of cells with a more epithelial appearance (Figure S1I). In agreement with their morphology, inhibition of EZH2 activity induced the expression of mesenchymal genes (Figure 1J and S1J) that were H3K27me3-positive prior to EZH2 inhibition (Figure S1K). The precise set of EMT genes induced varied in the three analysed cell lines (Figure S1M), in fitting with previous observation indicating EMT can be induced through different combination of EMT genes (Cook & Vanderhyden, 2020). Consistently, different combinations of mesenchymal TFs were activated in the three different cell lines (Figure S1L). Of note, *SNAI2* was gradually upregulated after 12 days of EZH2i treatment in exponentially growing MCF-7 cells (Figure S1D, compare day six and day twelve). After eighteen days of EZH2i-treatment, *SNAI1, TWSIT1* and *ZEB1* were also over-expressed. This suggests that persistent absence of H3K27me3 facilitates a time-dependent gradual activation of the EMT program and transition into the mesenchymal state.

To confirm that activation of mesenchymal genes upon EZH2i treatment we treated MCF-7 cells with another highly specific inhibitor of EZH2 that is approved for cancer treatment by the US Food and Drug Administration (EPZ-6438, tazemetostat) (Knutson *et al*, 2012), and measure its impact on gene expression by mRNA-seq. In keeping with our findings using EZH2i, treatment with EPZ-6438 inhibited H3K27me3 deposition (Figure S1N), induced the transcriptional activation of EZH2i-responsive genes (Figure S1O) and activated the mesenchymal gene expression program (Figure S1P). Overall, we concluded that H3K27me3 deposition through EZH2 is required to maintain the transcriptional repression of mesenchymal genes and favour the residence of breast carcinoma cells in an epithelial state.

### EZH2 is required to repress mesenchymal genes during TGF-β-dependent MET in breast cancer cells

To examine whether EZH2 regulates transitions between the epithelial and mesenchymal states of breast cancer cells we setup an in vitro system to study the dynamics of EMT and its reverse process (mesenchymal to epithelial transition, MET) (Figure S2A). Treatment of MCF-7 cells with 10 ng/ml transforming growth factor beta (TGF-β) and 50 ng/ml epidermal growth factor (EGF) during six days induced the transcription of mesenchymal markers (*N-CADHERIN, NRP2, TWIST1 and SNAI2*) and the downregulation of the epithelial marker *E-CADHERIN* (Figure S2B). Withdrawal of TGF-β and EGF from the culture media during six additional days led to the reversion of transcriptional changes: downregulation of the expression of mesenchymal genes (*N-CADHERIN, NRP2, TWIST1 and SNAI2*) coupled to activation of the epithelial marker E-CADHERIN (Figure S2B). Importantly, mRNA-seq analyses demonstrated that TGF-β stimulation induced a wide reversible reorganization of the transcriptome that involved 1108 genes (Figure S2C). This included the reversible activation of 750 genes enriched in EMT, cell migration and mesenchyme (cluster II, Figure S2C), as well as 358 reversibly repressed genes that included factors involved in estrogen receptor signalling (cluster I, Figure S2C). Importantly, changes in the expression of epithelial and mesenchymal genes were accompanied by expected functional changes in cell mobility in cultured wound healing assays (Figure S2D). We concluded that transient stimulation with TGF-β and EGF is a valid system to study EMT-MET in vitro in MCF-7 cells.

To test whether inhibition of EZH2 alters EMT-MET we compared the behaviour of the 1108 reversible genes identified in Figure S2C during transient TGF-β stimulation in the presence or absence of EZH2i and H3K27me3 marking (Figure 2A and 2B). Inhibition of EZH2 enhanced the transcriptional induction of 376 genes during EMT (cluster I, Figure 1C), and hindered the repression of 280 genes during MET (cluster II and cluster III, Figure 2C). This set of 461 genes were highly enriched for genes involved in EMT, TGF-ß stimulation, cell migration and mesenchyme (Figure 2D), indicating that EZH2 methyltransferase activity is required for reorganization of epithelial-mesenchymal gene expression programs during EMT-MET. In consonance, gene set enrichment analysis (GSEA) confirmed that MCF-7 cells treated with EZH2i displayed higher expression of mesenchymal genes than untreated cells (Figure 2E). Likewise, analysis of the expression of epithelial (*E-CADHERIN*) and mesenchymal (*N-CADHERIN, NRP2, TWIST1 and SNAI2*) genes in cells treated or untreated with EZH2i also supported that inhibition of EZH2 promotes activation of mesenchymal markers during EMT and hinders their repression during MET (Figure S2E). Expectedly, examination of the in vitro wound healing capacity of MCF-7 cells showed that treatment with EZH2i slightly increased the mobility of cells after six days of EMT (day six) and hindered the restoration of the more immobile epithelial phenotype upon MET (day twelve) (Figure 2F). Therefore, we settled that EZH2 is required to downmodulate the expression of mesenchymal genes during EMT-MET in breast cancer cells, and to allow efficient restoration of the epithelial state during MET.

**Figure 2.**
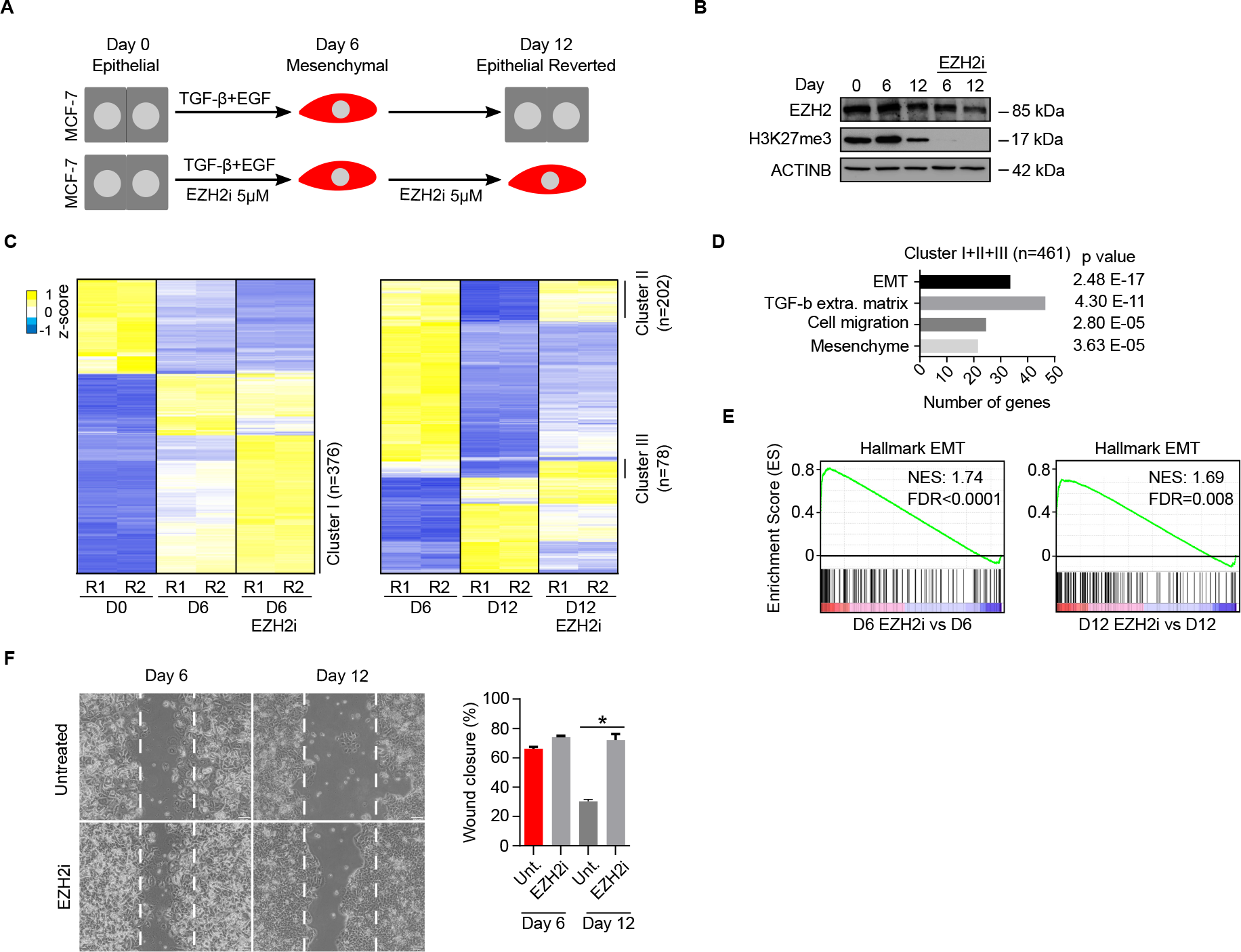
EZH2 is required to repress mesenchymal genes during TGF-β-dependent MET in MCF-7 cells. **(A)** Schematic diagram of the experimental conditions used to study the inhibition of EZH2 activity (EZH2i 5 μM) during reversible EMT-MET (TGF-β + EGF) in MCF-7 cells. **(B)** Western blot analysis of whole-cell extracts comparing the levels of EZH2 and H3K27me3 during the EMT-MET experiment described in figure 2A. ACTIN B provides a loading control. **(C)** Heatmap showing the expression of 1108 reversible genes (identified in figure S2C) during EMT-MET in two biological replicates (R1 and R2) at day 0, day 6 (upon EMT) and day 12 (upon MET), in the presence or absence of EZH2i, are shown. Genes that are differentially expressed due to the presence of EZH2i are labelled as clusters I, II and III. **(D)** Gene ontology analysis of genes identified in clusters I, II and III in figure 2C. **(E)** GSEA of EMT associated genes in cells treated with EZH2i, relative to untreated, at day 6 (left) or day 12 (right). Normalized enrichment score (NES) and statistically significant false discovery rate (FDR<0.25) are indicated. **(F)** Brightfield images and histogram analysing the effect of EZH2i treatment in wound healing closure after 72 hours, in cells corresponding to day 6 or 12 during the EMT-MET described in figure 2A. Mean and SEM of 3 experiments are shown. Asterisks indicate statistical significance using a Mann-Whitney test (* p<0.05).

### Expression of *EZH2* inversely correlates with the expression of mesenchymal genes in human breast tumours

To examine whether EZH2 represses mesenchymal genes in breast cancer cells in vivo, we used genome-wide gene expression datasets of 1904 resected breast cancer tumours available at the Molecular Taxonomy of Breast Cancer International Consortium. Importantly, we found that the levels of *EZH2* mRNA inversely correlate with the expression of EZH2 target genes (1126 out of the 1141 genes identified in figure 1A were analysable in these datasets) (Figure 3A). Negative correlation was more accused for the group of 197 bivalent genes that were induced upon treatment with EZH2i (Figure 3A). As expected, no significant correlation was found for a group of randomly selected control genes (Figure 3A). The expression values of individual mesenchymal genes such as *SNAI2* also displayed the expected negative correlation (Figure 3B). Therefore, these analyses support that EZH2 functions as a transcriptional repressor of the mesenchymal gene expression program in human breast cancer tumours. In agreement with previous reports (Adibfar *et al*, 2021; Jang *et al*, 2016), high expression of *EZH2* was associated with poor survival probability in our patient cohort (median survival, high: 132.3 months, low: 172.9 months) (Figure 3C). Overall, we concluded that augmented expression of EZH2 is associated to reduced expression of EZH2-target mesenchymal genes in breast cancer tumours and that high levels of EZH2 expression are associated with poor survival.

**Figure 3.**
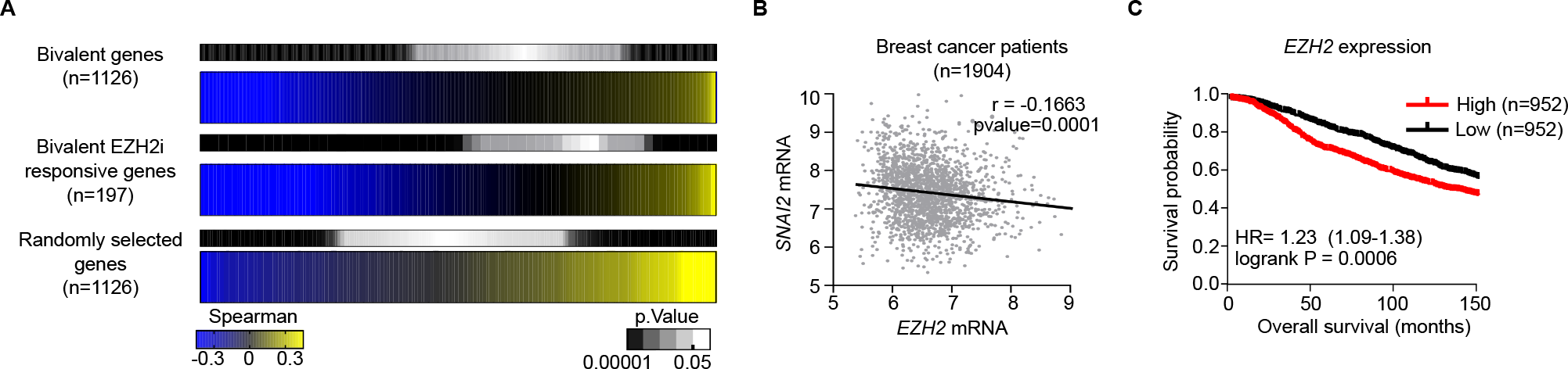
EZH2 level inversely correlates with the expression of mesenchymal genes, and it is associated with poor prognosis of breast cancer patients. **(A)** Heatmaps of Spearman’s correlation between the mRNA of *EZH2* and different subsets of genes in 1904 samples from breast cancer tumours. Bivalent genes include 1126 bivalent genes that were identified in figure 1A. Bivalent EZH2i responsive genes include 197 genes that are H3K27me3/H3K4me3-positive (figure 1A) and are included in cluster II in figure 1F. The pattern of a set of 1126 randomly selected genes is shown for comparison purposes. **(B)** Spearman’s correlation of *EZH2* and *SNAI2* mRNA in 1904 patients of breast cancer. Each dot represents expression values of one breast tumour sample. **(C)** Kaplan-Meier plots showing survival probability of 1904 breast cancer patients depending on the level of expression of *EZH2* mRNA.

## Discussion

Metastasis causes around 90% of cancer-associated mortality and therefore understanding the mechanisms underlying metastatic dissemination is crucial to develop more effective therapies in cancer (Lambert *et al*., 2017). Metastatic dissemination relies on dynamic changes in carcinoma cell state that occur during epithelial to mesenchymal reversible transitions and therefore EMT-MET has emerged as a key druggable pathway in cancer intervention (Dongre & Weinberg, 2019; Lambert *et al*., 2017). Our study reveals that the PRC2 catalytic subunit EZH2 coordinates the repression of the mesenchymal gene expression program in breast cancer cells, facilitating MET upon TGF-β stimulation decay. Because MET is required for efficient tumour colonization (Dongre & Weinberg, 2019; Lambert *et al*., 2017), our findings provide an explanation as to why EZH2-deficient breast cancer cells display reduced capacity to colonize new organs and form metastasis in mice models and patient-derived xenografts (Hirukawa *et al*., 2018; Moore *et al*., 2013; Yomtoubian *et al*., 2020; Zhang *et al*., 2022a), as well as to the oncogenic behaviour of *EZH2* as a marker of poor prognosis in breast cancer patients (this study and (Alford *et al*, 2012; Jang *et al*., 2016; Kleer *et al*, 2003)). Importantly, our discovery that EZH2 coordinates EMT-MET is in consonance with previous reports studying lung cancer cells where it has been shown that the loss of function of EZH2 hinders repression of mesenchymal genes during MET (Gallardo *et al*., 2022), and reduces tumour colonization capacity in mouse models (Gallardo *et al*., 2022; Serresi *et al*., 2016; Serresi *et al*., 2018). Counterintuitively, it has been recently reported that loss of function of a non-catalytic PRC2 subunit (EED) in experimentally transformed human mammary epithelial cells (HMLER) leads to increased metastasis in mouse xenograft experiments (Zhang *et al*., 2022b). This apparent discrepancy might rely on variations of EZH2 activity in the different loss of function systems: while in our experiments and previous reports in breast (Hirukawa *et al*., 2018; Yomtoubian *et al*., 2020) and lung (Gallardo *et al*., 2022; Serresi *et al*., 2016; Serresi *et al*., 2018) cancer the function of EZH2 was assayed in systems in which EZH2 protein level was reduced to undetectable levels, EED-depleted HMLER cells display only partial downregulation of EZH2 protein (Zhang *et al*., 2022b). We propose that low activity of EZH2 protein in EED-depleted HMLER cells promote the transition of breast cancer cells into a metastable mesenchymal state without fully impairing MET. This might explain the enhanced tumour colonization capacity observed in EED mutant cells, because latest reports indicate that cancer malignancy and disease progression rely on the ability of cancer cells to reside in intermediate metastable states within the epithelial – mesenchymal spectrum, rather than in extreme epithelial or mesenchymal states (Lüönd *et al*, 2021; Pastushenko *et al*, 2018; Simeonov *et al*, 2021). Overall, this study establishes a molecular framework that brings together previous reports in breast (Hirukawa *et al*., 2018; Moore *et al*., 2013; Yomtoubian *et al*., 2020; Zhang *et al*., 2022a; Zhang *et al*., 2022b) and lung (Gallardo *et al*., 2022; Serresi *et al*., 2016; Serresi *et al*., 2018) cancer cells, and supports a cell-of-origin-independent role of PRC2 as a direct modulator of the mesenchymal gene expression program during EMT-MET in human carcinomas.

## Methods

### Breast cancer cell lines culture conditions

The MCF-7, MDA-MB-231 and SKBR3 cell lines were kindly provided by the labs of Mª Jose Serrano and Dr. Juan Antonio Marchal (University of Granada, Spain). Cells were grown at 5% CO2 and 37 ºC in DMEM high glucose media supplemented with 10% heat inactivated fetal bovine serum (FBS) (Gibco), penicillin/streptomycin (Gibco), L-glutamine (Gibco) and 2-mercaptoethanol (Gibco). Detailed information about cell lines used is provided in Table S2.

### Induction of in vitro EMT-MET by TGF-β and EGF stimulation in MCF-7 cells

Epithelial MCF-7 cells were plated at a density of 10.000 cells/cm^2^ and treated with 10 ng/mL TGF-β (Prepotech) and 50 ng/mL EGF (Prepotech) for 6 days to induce transition into a mesenchymal state (EMT). Thereafter, both cytokines were removed from the culture media and cells were grown for 6 additional days to allow reversion to the epithelial state (MET). Cells were trypsinized, counted and replated at initial density in fresh media every 48h.

### Treatment of breast cancer cell lines with EZH2 inhibitors

EZH2 was inhibited using 5 μM GSK126 (A3446, APExBIO). Treatments with EZH2 inhibitor for 12 days were carried out by refreshing GSK126 every two days as MCF-7 cells were split to allow exponential cell growth. Likewise, during EMT-MET experiments, GSK126 was refreshed every two days as cells were diluted to the corresponding density (10 000 cells/cm^2^). In colony forming assays, cells were pre-treated with GSK126 for 4 days to reduce the global H3K27me3 level before plating the cells. MCF-7, SKBR3 and MDA-MB-231 cells were seeded at a density of 200 cells/cm^2^ to allow formation of colonies after 14 days. Media was refreshed every 5 days including GSK126 in the treated culture. In growth curves, cell viability and wound healing assays, cells were pre-treated with GSK126 for 4 days to reduce global H3K27me3 levels before the start of the experiment. Cells were maintained in the presence of GSK126 during the experiment as detailed below.

In experiments where EZH2 was inhibited using EPZ-6438, MCF-7 cells were exposed to 2 μM of EPZ-6438 (S7128, Deltaclon) for 12 days, and the inhibitor was refreshed every two days as MCF-7 were split to allow exponential cell growth.

### Growth curve, cell viability and in vitro wound healing assays

To perform growth curves, cells were plated at density of 10.000 cells/cm^2^ and diluted before confluence to the initial density. Accumulative growth was calculated by applying the dilution factor used. Inhibition of EZH2 was carried out by pre-treating cells with GSK126 for four days and refreshing the inhibitor in every cell dilution.

To analyse cell survival cells, were pre-treated with GSK126 for 4 days, plated in 96 well plate at a density of 3000 cells per well and allowed to grow. After 48h, cell media was replaced by fresh media containing 0.1mM of resazurin. The plate was incubated for 4h at 37ºC and fluorescence at 585 nm wavelength was measured. Same procedure was applied to GSK126 untreated control cells. Survival was estimated as the ratio of the signal measured in treated cells relative to untreated control.

In cultured wound healing assays MCF-7 untreated or pre-treated for 4 days with GKS126 were grown to 100% confluence. A 1000 μL sterile pipette tip was used to produce a scratch in the monolayer of cells. Standard cell media was change to media containing 1% of FBS and supplemented or not with GSK126. Cells were allowed to close the wound for 48 hours. The area of the wound was imaged every 24 hours using a widefield microscope and quantified using Image J software. The area of the wound at each time point was normalized to the area of the wound at time zero.

### RT-qPCR

RNA was isolated using Trizol reagent (Thermofisher), reverse transcribed using RevertAid Frist Strand cDNA synthesis kit (Thermofisher) and analysed by SYBRG real-time PCR using GoTaq qPCR Master Mix (Promega). Primers used are provided in supplementary Table S2.

### Western blot

Western blots of whole cell extracts, or histone preparations were carried out using standard procedures as previously described (Gallardo *et al*, 2020). The following primary antibodies were used: rabbit anti-EZH2 (Diagenode), mouse anti-H3K27me3 (Active Motif), rabbit anti-SNAI2 (CST), mouse anti-E-CADHERIN (BD), mouse anti-ACTIN B (Sigma-Aldrich), rabbit anti-ACTIN B (Cell signalling). Secondary species-specific antibodies conjugated to horseradish peroxidase were used: anti-rabbit-HRP (GE-Healthcare), anti-mouse-HRP (GE-Healthcare) and anti-goat-HRP (Abcam). Clarity Western ECL reagents (Bio-Rad) was used for detection. More information about antibodies used is provided in Table S2.

### ChIP sequencing analysis

Public ChIP-seq datasets of H3K27me3 and H3K4me3 performed in MCF-7, MDA-MB-231 and SKBR3 (Table S2) were analysed as follows. Alignment of the sequence reads was done using Bowtie2 (Langmead & Salzberg, 2012) and human genome hg19 were used for mapping. Unmapped and multimapped reads were filtered out with SAMTools (Li *et al*, 2009) and SamBamba (Tarasov *et al*, 2015) to keep only uniquely aligned sequences. BigWigs were generated after normalizing with their input signal using the BamCompare function of the deepTools suite package (Richter *et al*, 2016). Peak calling was performed with MACS version 3 (Zhang *et al*, 2008) using the input as background for normalization. Peaks with q ≤ 0.05 values were considered significant. Significant peaks were annotated with Homer software (Heinz *et al*, 2010) by defining promoter regions as ±2 kb from the start of the transcription start site (TSS). The coverage of the samples around the TSS was performed with the Bioconductor package coverageView for a genomic window of ± 4kb using a bin size of 10 bp. R version 4.2.2 and RStudio version 2022.7.1.554 was used. BigWigs were generated using the deepTools suite (Ramírez *et al*, 2014) and reads per million (RPM) were used to represent the ChIP-seq signal.

### mRNA sequencing analysis

Total RNA was isolated using Trizol reagent (Thermofisher) or Rnease mini kit (Quiagen). Libraries and sequencing were performed at BGI Genomics. Strand specific mRNA-seq libraries were generated using 200 ng of total RNA and the DNBSEQ library construction protocol. Libraries were sequenced using DNBSEQ high-throughput platform sequencing technology. 20million (150 bp paired-end) reads were obtained for each condition.

RNA sequencing data was analysed using the miARma-Seq pipeline (Andrés-León *et al*, 2016). First, quality control of reads was performed using FastQC software (Andrews, 2010). Reads were aligned using STAR 2.5.3a against the reference human genome hg19 (GENCODE assembly GRCh37.p13). To obtain expression values featureCounts 2.0.6 (Liao *et al*, 2014) was used. Reference gene annotations were obtained from GENCODE assembly mentioned above. Normalization of gene expression values was obtained applying the trimmed mean values method (TMM) (Robinson & Oshlack, 2010) using the NOISeq package (Tarazona *et al*, 2015). Differential gene expression analysis was performed using DESeq2 package (Love *et al*, 2014). GSEA (Subramanian *et al*, 2005) was performed against the set “Hallkmark Epithelial to Mesenchymal Transition” from the Molecular Signatures Database (MSigDB) (Liberzon *et al*, 2015).

### Bioinformatic analysis of breast cancer tumour samples

Information of 1904 breast cancer tumour samples from the Breast Cancer METABRIC dataset were analysed using CBioportal tools. Kaplan Meier survival analysis and the correlation analyses between mRNA levels of EZH2 of selected target genes were performed.

### Statistical analyses

Statistical significance (p<0.05) was determined by applying a two-tailed non-parametric Mann-Whitney test. Spearman’s correlation coefficient was calculated to measure correlation among variables. All analyses were performed with GraphPad prism 9 and/or R or Rstudio.

### Data access

Datasets are available at GEO-NCBI with accession number GSE247138. Temporal password will be eliminated upon publication acceptance.

## Author contributions

DL designed and conceptualized the study. AG, LLO and DL designed experiments. AG, LLO, ASP and AM performed experiments. EBR and IFR performed bioinformatic analyses. LLO and ASP provided resources and supervision of experiments. AG and DL wrote the manuscript. All authors revised the manuscript. DL obtained funding and supervised research.

## Funding

The Landeira lab is supported by the Spanish Ministry of Science and Innovation (PID2019-108108-100, EUR2021-122005), the Instituto de Salud Carlos III (IHRC22/00007), the Andalusian regional government (PC-0246-2017, PIER-0211-2019, PY20_00681) and the University of Granada (A-BIO-6-UGR20) grants. Efres Belmonte-Reche is funded by a Maria Zambrano fellowship. Lourdes Lopez-Onieva is supported by the “Plan-Propio UGR” tenure track program.

## Acknowledgements

We are very grateful to Dr. Maria Jose Serrano, Dr. Sergio Granados-Principal and Dr. Juan Antonio Marchal for providing cell lines. We thank core facilities at GENYO for excellent technical support.

## Competing interests

The authors declare no competing interests.

---

**Figure S1.**
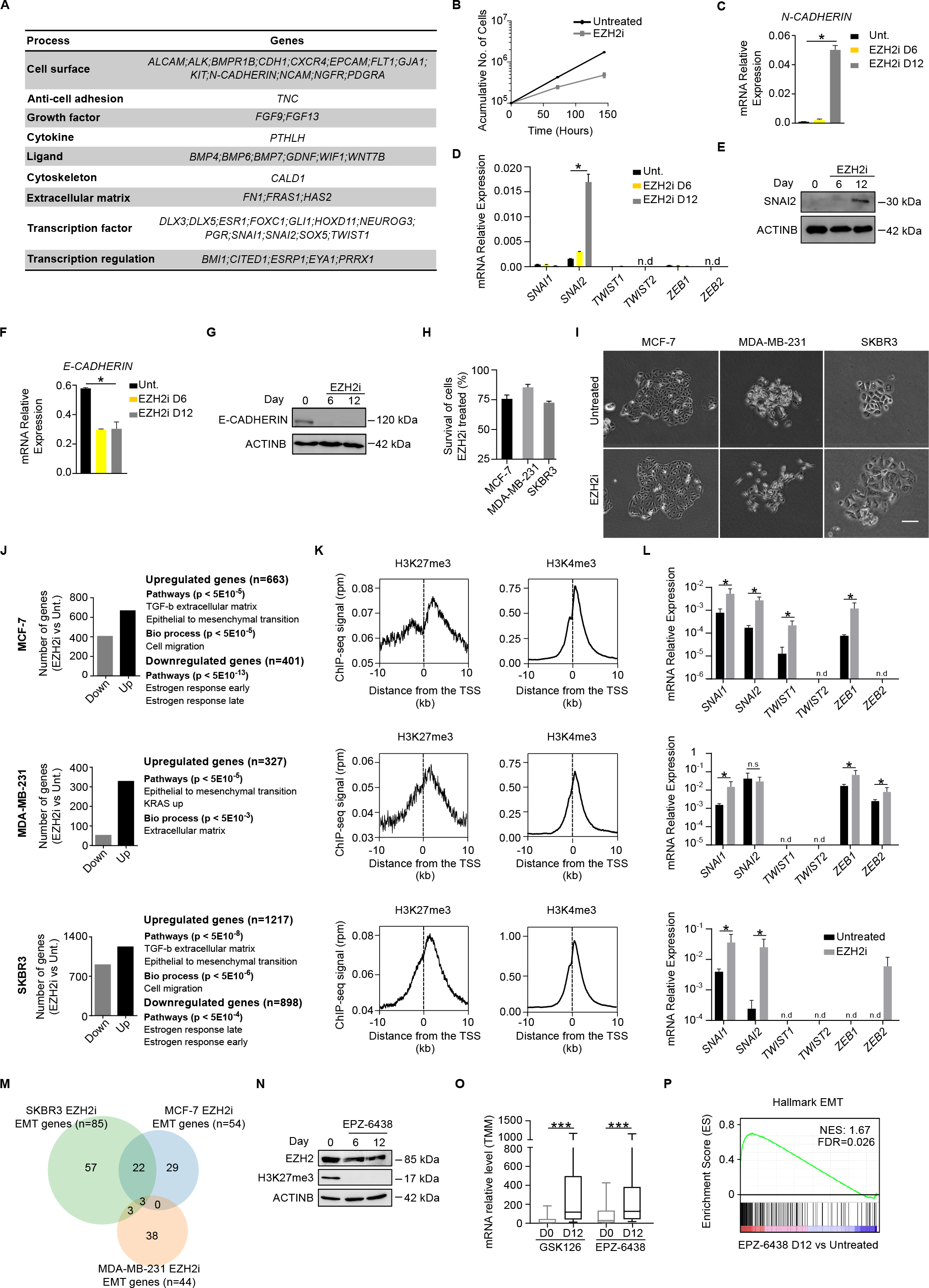
Treatment with EPZ-6438 induces the activation of mesenchymal genes in MCF-7 cells. **(A)** Table showing bivalent genes classified related to mesenchyme identified in figure 1C. (B) Growth curve of MCF-7 cells treated with EZH2i and untreated control cells. **(C)** Analysis of *N-CADHERIN* mRNA expression by RT-qPCR during EZH2i treatment in MCF-7cells. Expression is calculated relative to housekeeping genes *GAPDH* and *ACTINB*. **(D)** Analysis of EMT-TFs mRNA expression by RT-qPCR during EZH2i treatment in MCF-7cells. Expression is calculated relative to housekeeping genes *GAPDH* and *ACTINB*. **(E)** Western blot analysis of SNAI2 in whole-cell extracts from EZH2i-treated MCF-7 cells. ACTIN B was used as a loading control. **(F)** Histogram show the level of *E-CADHERIN* mRNA expression measured by RT-qPCR during EZH2i treatment in MCF-7 cells. Expression is calculated relative to housekeeping genes *GAPDH* and *ACTINB*. **(G)**Western blot analysis of the level of E-CADHERIN in whole-cell extracts from EZH2i-treated MCF-7 cells. ACTIN B was used as a loading control. **(H)** Histogram showing cell survival of indicated cells lines treated with EZH2i relative to untreated control after resazurin staining and fluorescence quantification. **(I)** Brightfield images of breast cancer cell lines plated at low density in the presence of EZH2i for 14 days. **(J)** Histogram showing the number of genes downregulated (grey bar) or upregulated (black bar) (FC > 2, p < 0.05) in indicated cell lines after plating cells at low density in the presence of EZH2i for 14 days. Terms enriched in gene ontology analyses are shown. **(K)** Average enrichment ChIP-seq signal of H3K27me3 and H3K4me3 around the TSS of genes induced upon treatment with EZH2i in indicated cell lines. **(L)** Histograms showing the level of expression of mesenchymal TFs in indicated cell lines treated or untreated with EZH2i conditions for 18 days. **(M)** Venn diagram comparing sets of EMT genes induced after treatment with EZH2i for 14 days. **(N)** Western blot analysis of whole-cell extracts comparing the levels of EZH2 and H3K27me3 during treatment of MCF-7 cells with 2 μM EPZ-6438 for 12 days. ACTINB provides a loading control. **(o)** Boxplot comparing mRNA expression of 573 genes responsive to EZH2 inhibition (cluster II in Figure 1F) by RNA-seq in MCF-7 cells treated or untreated for twelve days with EZH2 inhibitors GSK126 and EPZ-6438. Asterisks indicate statistical significance using a Mann-Whitney test (*** p<0.001). **(P)** GSEA of EMT associated genes in MCF-7 cells treated with EZH2 inhibitor EPZ-6438 for 12 days. Normalized Enrichment Score (NES) and statistically significant false discovery rate (FDR<0.25) are indicated. Mean and SEM of 3 experiments are shown in C, D, F and M. Asterisks indicate statistical significance using a Mann-Whitney test (* p<0.05).

**Figure S2.**
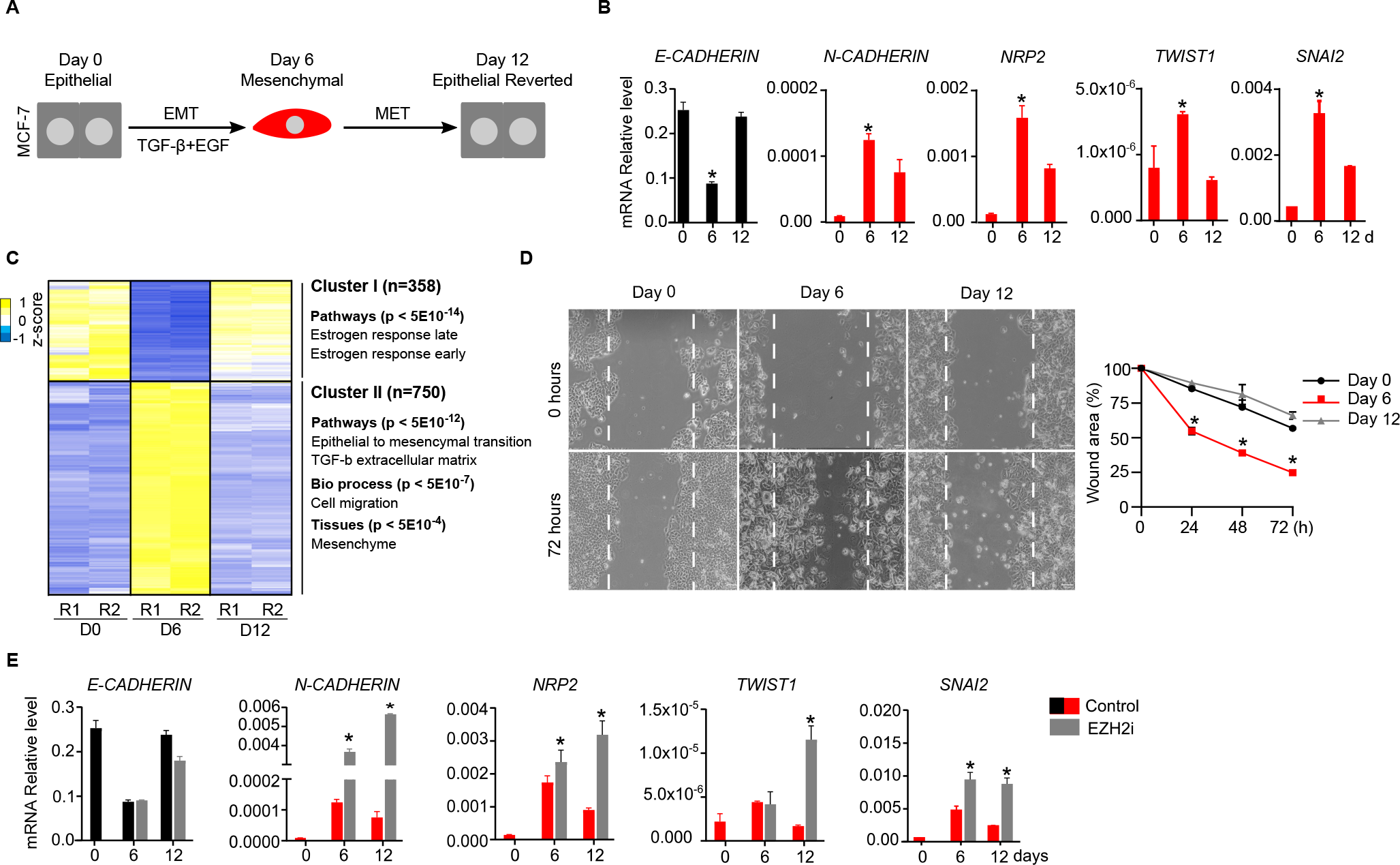
TGF-β-induced EMT is reversible in MCF-7 cells. **(A)** Scheme of the experimental design used to induce EMT-MET in MCF-7 cells. **(B)** Analysis of mRNA expression by RT-qPCR of indicated epithelial (black) or mesenchymal (red) genes during EMT-MET. Relative expression level against *GAPDH* and *ACTIN B* is shown. **(C)** Heatmap showing mRNA expression of 1108 genes that were reversibly regulated (FC > 2, p < 0.05) during EMT-MET. Two independent replicates (R1 and R2) at day 0 (E), day 6 (M) and day 12 (ER) are shown. Gene ontology analyses of genes in indicated clusters are shown. **(D)** Brightfield images and quantification plot comparing the wound healing capacity after 72 hours of cells obtained at day 0, 6 and 12 during EMT-MET. **(E)** RT-qPCR analysis showing mRNA level of indicated epithelial (black) and mesenchymal (red) genes during EMT-MET in the absence or presence of EZH2i. Expression level is calculated relative to *GAPDH* and *ACTIN B*. Mean and SEM of 3 experiments are shown in B, D and E. Asterisks indicate statistical significance using a Mann-Whitney test (* p<0.05).

## Notes

### Competing Interest Statement

The authors have declared no competing interest.

### Summary of Updates

mRNA-seq experiments supporting previous findings by RT-qPCR.

